# Large-scale Characterization of Drug Responses of Clinically Relevant Proteins in Cancer Cell Lines

**DOI:** 10.1101/2020.07.03.186908

**Authors:** Wei Zhao, Jun Li, Mei-Ju Chen, Zhenlin Ju, Nicole K. Nesser, Katie Johnson-Camacho, Christopher T. Boniface, Yancey Lawrence, Nupur T. Pande, Michael A. Davies, Meenhard Herlyn, Taru Muranen, Ioannis Zervantonakis, Erika Von Euw, Andre Schultz, Shwetha V. Kumar, Anil Korkut, Paul T. Spellman, Rehan Akbani, Dennis J. Slamon, Joe W. Gray, Joan S. Brugge, Yiling Lu, Gordon B. Mills, Han Liang

**Author notes:** Correspondence: H.L., (lead contact) and G.B.M. These authors contributed equally to this study (Wei Zhao, Jun Li).

## Abstract

Perturbation biology is a powerful approach to developing quantitative models of cellular behaviors and gaining mechanistic insights into disease development. In recent years, large-scale resources for phenotypic and mRNA responses of cancer cell lines to perturbations have been generated. However, similar large-scale protein response resources are not available, resulting in a critical knowledge gap for elucidating oncogenic mechanisms and developing effective cancer therapies. Here we generated and compiled perturbed expression profiles of ~210 clinically relevant proteins in >12,000 cancer cell-line samples in response to >150 drug compounds using reverse-phase protein arrays. We show that integrating protein response signals substantially increases the predictive power for drug sensitivity and aids in gaining insights into mechanisms of drug resistance. We build a systematic map of protein-drug connectivity and develop an open-access, user-friendly data portal for community use. Our study provides a valuable information resource for a broad range of quantitative modeling and biomedical applications.

**Highlights:**

- A large collection of cancer cell line protein responses to drug perturbations
- Perturbed protein responses greatly increase predictive power for drug sensitivity
- Build a systematic map of protein-drug connectivity based on response profiles
- Develop a user-friendly, interactive data portal for community use

## Introduction

Cancer is a highly heterogeneous disease encompassing many tissue types and diverse oncogenic drivers, with treatment responses that are often variable in distinct tumor contexts. Over the last decade, extensive efforts have been made to characterize the tremendous heterogeneity of human cancers at the molecular level (Berger et al., 2018; Hutter and Zenklusen, 2018; Jiang et al., 2019; Liu et al., 2018; Taylor et al., 2018). A real challenge in cancer research, however, is to obtain a systematic understanding of causality and mechanisms underlying the behaviors of cancer cells with the eventual goal of improving patient outcomes (Wise and Solit, 2019). To address this challenge, perturbation experiments provide a powerful approach in which cells are modulated by perturbagens, and downstream consequences are monitored (Korkut et al., 2015; Molinelli et al., 2013; Ng et al., 2018). The longitudinal data thus obtained provide considerably greater information content of both the basal biological network wiring and its associated changes under stress, thereby leading to a deeper understanding of mechanisms underlying cell survival under stress. Recently, large-scale compendia of the phenotypic and cellular effects of perturbed cancer cell lines have been established. For example, large-scale pharmacologic perturbation studies, cell viability measurement upon different drug treatments across many cell lines, have been published (Barretina et al., 2012; Basu et al., 2013; Garnett et al., 2012; Iorio et al., 2016); several studies have built genome-wide “cancer dependency” maps across a large number of cell lines using loss-of-function siRNA, shRNA, or CRISPR/cas9 screens (McDonald et al., 2017; Tsherniak et al., 2017); a “connectivity map” of profiled mRNA responses of cancer cell lines to diverse perturbations using an efficient, robust RNA measurement platform, L1000 has been developed (Subramanian et al., 2017). These studies provide valuable resources for gaining a systems-level understanding of cancer mechanisms and phenotypes. However, similar large-scale resources for analysis and integration of protein responses of perturbed cancer cell lines have yet to be established. This knowledge gap is even more striking, considering that proteins comprise the basic functional units in biological processes and represent the major targets for cancer therapy.

To fill this gap, we generated and compiled a large compendium of perturbed protein expression profiles of cancer cell lines in response to a diverse array of clinically relevant drugs using reverse-phase protein arrays (RPPAs). As a quantitative antibody-based assay, RPPA can assess a large number of protein markers in many samples in a cost-effective, sensitive, and reproducible manner (Hennessy et al., 2010; Nishizuka et al., 2003; Tibes et al., 2006). We have applied this technology to quantify protein expression levels of large patient cohorts (e.g., The Cancer Genome Atlas) (Akbani et al., 2014; Zhang et al., 2017) and cancer cell lines (e.g., MD Anderson Cell Line project and Cancer Cell Line Encyclopedia) (Ghandi et al., 2019; Li et al., 2017). The current antibody repertoire covers key oncogenic pathways such as PI3K/AKT, RAS/MAPK, Src/FAK, TGFβ/SMAD, JAK/STAT, DNA damage repair, Hippo, cell cycle, apoptosis, histone modification, and immune-oncology. Compared with proteome-wide mass spectrometry approaches, our RPPA-based approach has several advantages. First, although the number of protein markers in RPPA readout is much smaller (~200), this highly select protein set is enriched in therapeutic targets and biomarkers, thereby greatly increasing the ability to generate clinically relevant hypotheses and make translational impacts. Statistically speaking, this more focused assessment also substantially reduces the burden of multiple testing, a major challenge in identifying significant hits from unbiased proteomic searches. Second, one RPPA slide can measure up to 1,000 samples simultaneously. Thus, the high-throughput and cost-effectiveness make RPPA a practical platform for assessing a large number of samples (e.g., >10,000), which is simply not feasible for alternative proteomic approaches. Third, protein-level responses, particularly changes in post-translational modifications, more likely reflect how cancer cells rewire their signaling pathways to adapt and survive a specific drug treatment, as most targeted therapies act by modulating protein phosphorylation and activity. The superior ability of RPPA to quantify some key post-translationally modified proteins has the potential to capture such adaptive responses and can provide stronger predictors of therapy response or resistance mechanisms (Mertins et al., 2014). Indeed, our recent studies have demonstrated the value of RPPA-based adaptive responses in the rational design of combination therapies (Fang et al., 2019; Iavarone et al., 2019; Korkut et al., 2015; Krepler et al., 2017; Krepler et al., 2016; Kwong et al., 2015; Molinelli et al., 2013; Muranen et al., 2012; Sun et al., 2017; Sun et al., 2018), with several of these translated to the clinic (NCT01623349, NCT03586661, NCT02208375, NCT02338622, NCT03162627; NCT03579316, NCT03565991, NCT03801369, NCT03544125) with patient benefit.

## Results

### A large, high-quality collection of perturbed protein expression profiles of cancer cell lines

To generate a high-quality resource of perturbed protein responses, we measured RPPA-based protein expression profiles of cancer cell lines in response to >150 preclinical and clinical therapeutics (often across multiple time points), generated normalized RPPA data (including baseline level p0 and post-treatment level p1) and protein response to perturbation (Δp = p1 – p0) profiles using a standardized data processing pipeline, and made the data public through a user-friendly data portal (**Figure 1A**). In total, this compendium contains RPPA profiles (~210 total and phosphorylated protein markers) of 15,867 samples (12,222 drug-treated samples and 3,647 control samples). The cancer cell lines come from several lineages, including breast, ovarian, uterus, skin, blood, and prostate; and the drug compounds target a broad range of cancer-related processes, including PI3K/mTOR signaling, ERK/MAPK signaling, RTK signaling, EGFR signaling, TP53 pathway, genome integrity, cell cycle, antipsychotic drugs, and chromatin remodeling (**Figure 1B**). Due to time and cost constraints and the clinical relevance of different drugs, instead of profiling all possible perturbations across all cell-line and drug combinations, we took a more pragmatic approach in which some cell lineages and drug groups were more frequently profiled but still represent an extensive survey of drug perturbations (**Figure 1B)**. Our sample set is highly enriched in responses from a subset of common, well-characterized cancer cell lines that have rich molecular profiling and drug response data in public resources (**Figure 1C**). For example, >1,500 drug-treated samples were from MCF7, and >250 drug-treated samples were from BT20, SKBR3, MDA-MB-468, BT549, UACC812, BT474, SKOV3, and HCC1954 (**Figure S1A**). For drug treatment, 86.2% of the samples were treated with monotherapy, and ~1,700 samples were treated with double or triple-drug combinations (**Figure S1B**). Among the drug compounds used, 23 compounds have >150 treated samples, with lapatinib (485 samples, HER2 inhibitor), GSK690693 (453 samples, AKT inhibitor), and AZD8055 (424 samples, mTOR inhibitor) being the top three (**Figure S1C**). Importantly, for many of the therapeutic targets, we profiled multiple targeting agents, including those that target different members of the same pathway, to increase our ability to identify on-target activity. To assess overall data quality, we compared protein response (Δp) correlations of technical replicate samples (n = 2,771 pairs) to those of randomly selected sample pairs. We found that replicate samples showed much higher correlations across protein markers (mean R = 0.87) than random pairs (mean R = 0.059) (**Figure 1D**), indicating high reproducibility of our RPPA data.

**Figure 1.**
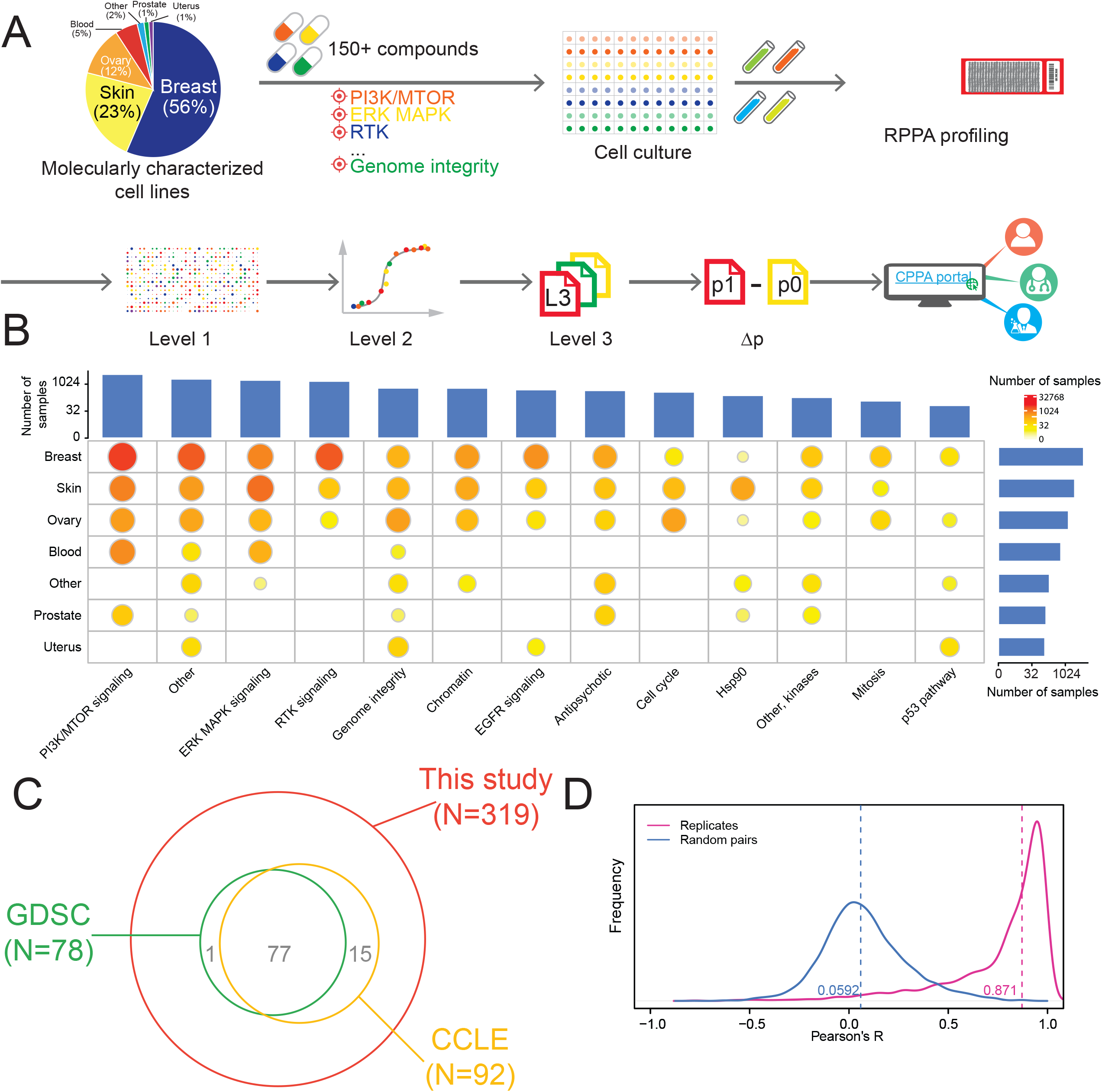
Summary of the perturbed RPPA profiling data in this study. Overview of the RPPA profiling experiments and data processing of cell line perturbations. The pie chart shows the lineage distribution of cancer cell lines (n = 319) profiled. (B) The distribution of drug-treated samples by cell lineages and drug groups. The bar plots show the numbers of samples profiled for each lineage or drug group, and the size of the circle is proportional to the number of samples profiled for each lineage-drug combination. (C) The circles show the number of cell lines used in this study that were profiled by GDSC and CCLE. (D) Reproducibility of perturbed RPPA data based on technical replicates (n = 2,771 pairs). See also Figure S1.

To further confirm the quality of the RPPA data output, we sought to validate our protein response data using independent mRNA response data from the connectivity map (Subramanian et al., 2017). Since this analysis is for different molecules (protein vs. RNA) and across different platforms (RPPA vs. L1000), we employed the Goodman-Kruskal’s gamma (ɣ) correlation to conduct a robust assessment. Based on the same cell lines perturbed by the same compounds (n = 46 unique cell-line-drug perturbations), we first converted the original continuous response scores into categorical response groups (i.e., upregulated, neutral, and downregulated) and then compared the mRNA-protein response concordance by calculating mRNA-protein response association and sample-sample association (**Figure 2A**). We observed that the matched mRNA-protein responses from the same condition were highly associated with each other (median ɣ = 0.63), which is significantly higher than that from the randomly shuffled background distribution (paired Student’s t-test, p = 5.8×10^−5^, **Figure 2B**). Then, we tested whether the sample-sample association inferred from the RPPA-based protein responses were preserved in the L1000-based mRNA responses. Among the significant sample-sample associations identified by either platform (FDR < 0.01), the RPPA-based ɣ scores showed a strong, positive correlation with the L1000-based ɣ scores (Pearson’s correlation, R = 0.65, p = 5×10^−6^). Further, categorized RPPA-based associations are highly consistent with L1000-based associations (Fisher’s exact test, p = 2.3×10^−3^). These cross-molecule, cross-platform, and cross-study comparisons strongly support a high quality of the protein response data.

**Figure 2.**
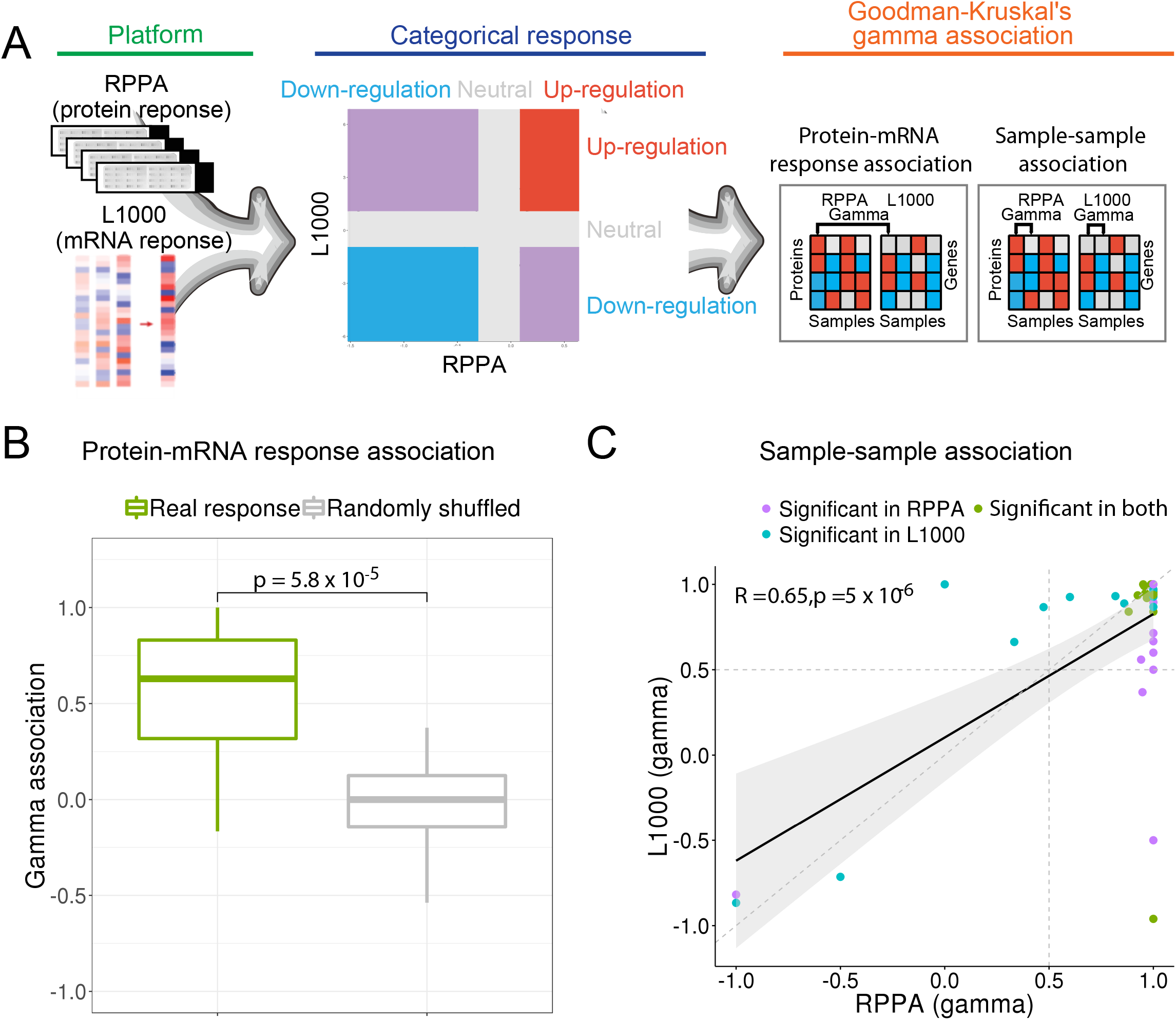
Comparison between the RPPA-based protein responses and the L1000-based mRNA responses. (A) Overview of the comparison method (see STAR Methods for details). (B) Boxplots of protein-mRNA response associations between the RPPA and L1000 platforms using the same perturbations (e.g., the same cell line and the same compound, n = 46). The gamma associations from the real responses (green box) were compared to those from the randomly shuffled background distribution (grey box). The P-value is based on a paired Student’s t-test. The vertical line in the box is the median, the bottom, and top of the box are the first and third quartiles, and the whiskers extend to 1.5 IQR of the lower quartile and the upper quartile, respectively. (C) Scatter plot showing the correlation of sample-sample gamma associations from the RPPA (x-axis) and L1000 (y-axis) platforms. Only significant data points (gamma associations) with FDR < 0.01 in either platform are shown. Pearson’s correlation coefficient and associated P-value are shown.

### Predictive power and mechanistic insights for drug sensitivity by protein responses

Our previous study demonstrated that RPPA-based baseline protein levels showed considerable predictive power for drug sensitivity in cancer cell lines (Li et al., 2017). To assess the predictive power of protein responses for drug sensitivity, we integrated our perturbed RPPA data and drug sensitivity data available in GDSC (Iorio et al., 2016) and identified seven drugs whose sensitivity and protein expression data were available in at least six different cell lines. Then, for each drug, we defined three types of protein markers that may be informative about drug sensitivity: (i) p0: the baseline expression of a protein shows a significant correlation with the sensitivity to the drug across cell lines (Pearson correlation, p < 0.05); (ii) Δp only: the protein response shows a significant correlation with drug sensitivity (Pearson correlation, p < 0.05); and (iii) Δp|p0: given p0, the protein response shows additional information content in predicting drug sensitivity (see STAR Methods). Across all the drugs, the numbers of Δp-informative (Δp only + Δp|p0) protein markers were significantly higher than those of p0-based markers (paired t-test, p = 1.38×10^−3^, n = 7 drugs, **Figure 3A**). We next focused on two representative drugs, pictilisib (PI3K inhibitor), and talazoparib (PARP inhibitor), with markedly different targets, for which a large number of cell lines have drug sensitivity data (Iorio et al., 2016; Seashore-Ludlow et al., 2015). We found that across cell lines, baseline proteins (p0) and protein response (Δp) showed distinct sets of proteins whose levels significantly correlated with drug sensitivity (Pearson correlation, p < 0.05), respectively; and there were additional proteins where Δp correlated with drug sensitivity when considering the information content of p0 (Δp|p0). When combining Δp and Δp|p0, the number of informative protein responses increased dramatically compared to baseline protein levels: pictilisib, from 17 to 27; and talazoparib, 33 to 52 (**Figure 3B, 3C**). Considering the potential noise of drug sensitivity data, we further validated this pattern, either using independent public drug sensitivity data or generating in-house drug sensitivity data (**Figure S2**). The correlations of baseline protein levels or protein responses with drug sensitivity between the different resources are highly correlated, despite the assessment of independent cell line sets (**Figure 3D, Figure S2**). These results not only further support the high quality of the RPPA expression data, but also suggest that changes in protein levels on therapeutic challenge provide substantial additional information content beyond that provided by baseline protein levels for predicting treatment responses.

**Figure 3.**
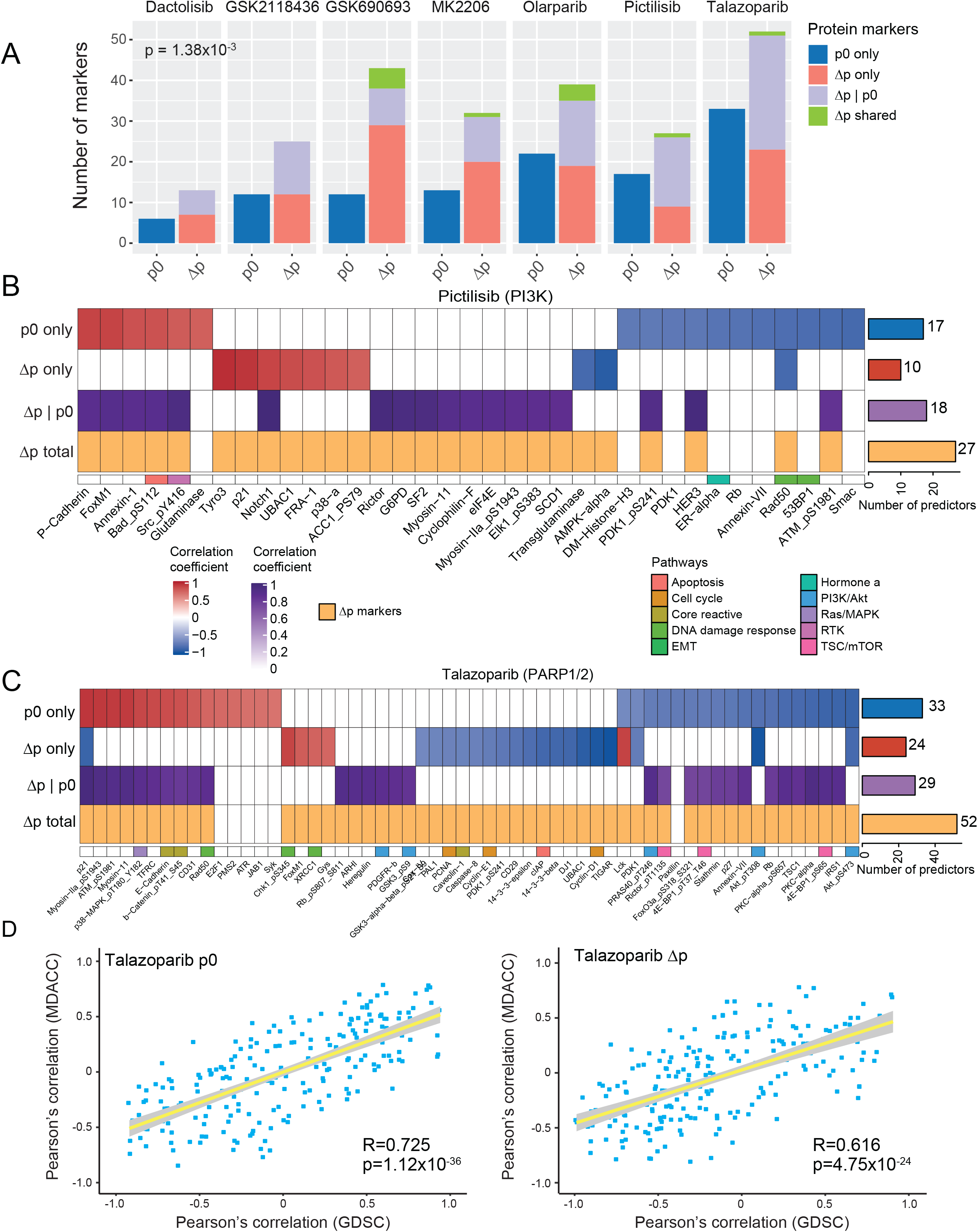
Comparison of the predictive power of protein markers for drug sensitivity. (A) Summary of predictive markers based on baseline protein expression (p0) and protein response (Δp) using drug response data from GDSC. Given a specific drug, three types of predictive markers were identified: (i) proteins whose p0 level is significantly correlated with drug sensitivity; (ii) proteins whose Δp level is significantly correlated with drug sensitivity; and (iii) proteins whose Δp level is significantly correlated with drug sensitivity given the p0 contribution. These protein markers were identified based on both Δp only, and Δp|p0, and are called Δp shared. (B, C) Heatmaps showing different types of protein markers for GDC0941/pictilisib (B) and BMN673/talazoparib (C). In each panel, the heatmap shows Pearson’s correlation coefficients of different protein levels with sensitivity to the drug of interest; the bar plot shows the number of predictive protein markers in each category. The protein markers of baseline (p0) and protein response (Δp) were selected at a significance level of p = 0.05 in univariate correlation. The joint protein markers, labeled as “Δp|p0,” represent linear regression models, including both p0 and Δp levels for a specific protein. The coefficients in the heatmap for joint markers indicate the correlation of the prediction value of the linear model with the real value of drug sensitivity. The proteins identified by Δp only or “Δp|p0” are marked by Δp total. (D) The scatter plots summarize the comparisons of Pearson’s correlation coefficients of drug sensitivity and baseline level (p0, left) as well as protein response (Δp, right) using two independent data sets. See also Figure S2.

To demonstrate how protein response could help elucidate drug resistance mechanisms and suggest therapy combinations, we focused on MEK inhibitors (MEKi), using cobimetinib as an illustration example and considering both baseline (p0) and protein response levels (Δp) (**Figure 4**). Cell lines were divided into MEKi-resistant (OVCAR432: RAS pathway WT, OVCAR3: RAS pathway WT, and OAW28: MAP2K4 mutant) and MEKi-sensitive (OV90: BRAF mutant, CAOV3: RAS pathway WT, ES2: BRAF and MEK mutant, OVCAR5: KRAS mutant, JHOM1: RAS pathway WT, and OVCAR8: KRAS mutant) based on response to multiple MEK inhibitors in our and publicly available data (CTRPv2 and GDSC). As expected, cell lines with aberrations in the RAS/MAPK pathway have a higher propensity to RAS/MAPK baseline pathway activity and sensitivity to MEKi, as indicated by low BIM and high EGFR, DUSP4, transglutaminase, pYB1, p90RSK, pMAPK, pMEK, and pJun (Pohl et al., 2005) (**Figure 4A**). There was also a suggestion that cell state and, particularly, decreased epithelial characteristics, or epithelial-mesenchymal transition (EMT) (low E-cadherin, beta-catenin, RAB25, ERalpha, GATA3, high EPPK1, N-cadherin, AXL, PAI-1, and fibronectin) were associated with sensitivity to MEKi. The EMT characteristics were likely mediated, at least in part, by effects of the RAS/MAPK pathway activation noted above (Shao et al., 2014).

**Figure 4.**
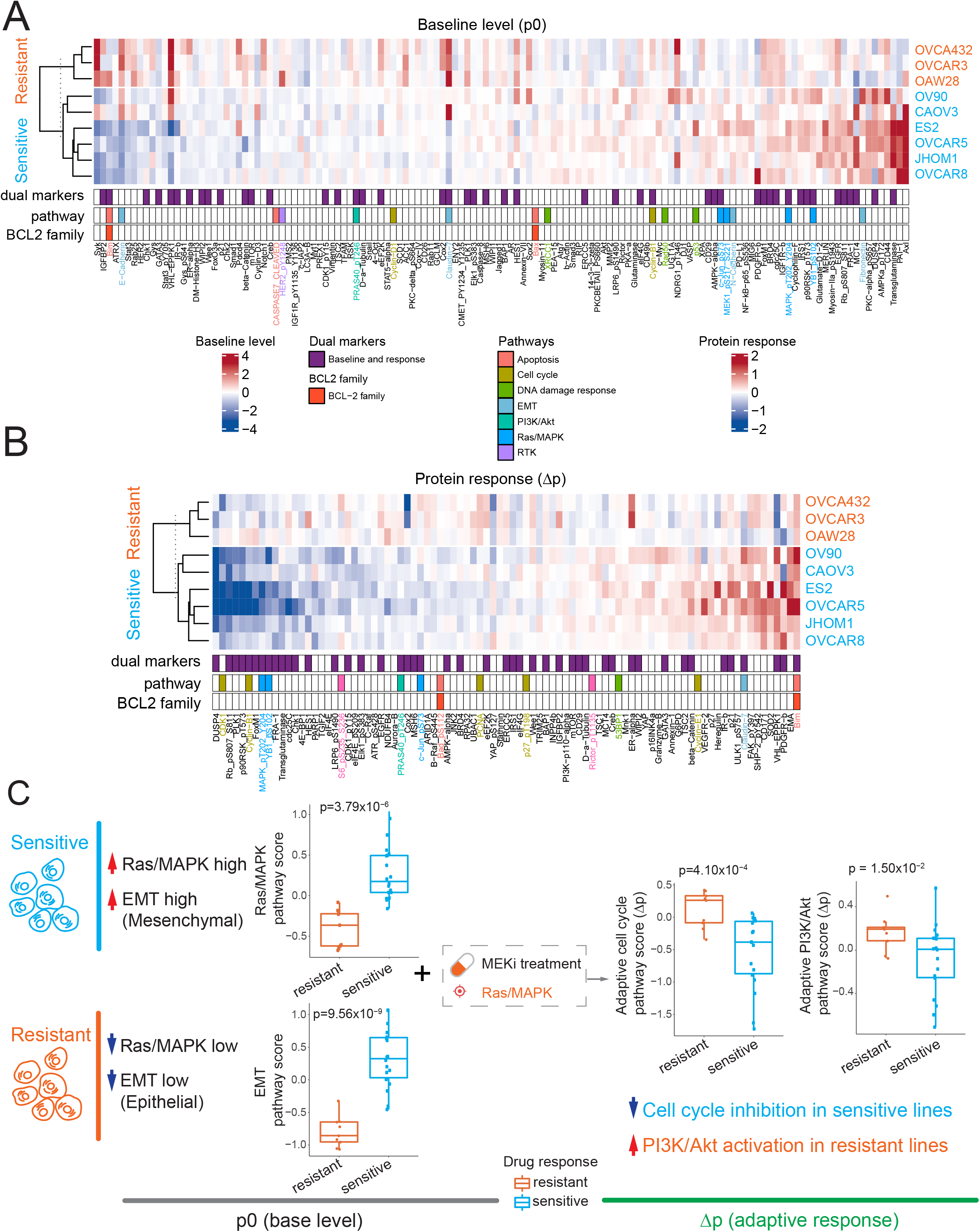
Differentially expressed protein markers between cobimetinib-sensitive and resistant cell lines. (A, B) Heatmaps showing baseline (A) and perturbed protein expressions (B) with significant differences between sensitive and resistant cell lines (q < 0.05). Each protein marker is annotated by whether it is a dual marker (i.e., significant both in p0 and Δp), BCL-2 family member, or belongs to a specific pathway. (C) Cartoon summary of baseline protein levels and adaptive protein responses to MEK inhibitors between the two cell groups. The difference of pathway scores between the two groups was assessed based on a Student’s t-test. See also Figure S3.

Adaptive responses (Δp) to cobimetinib demonstrated a greater dynamic range in terms of sensitivity and resistance to MEKi than baseline data. Sensitivity to cobimetinib was associated with evidence for a greater cobimetinib-induced decrease in RAS/MAPK pathway activity (decreased DUSP4, transglutaminase, FOXM1, p90RSK, pMAPK, pYB1, pS6, and pJun, and increased BIM), and decreased cell cycle progression (decreased pRB, cyclinB1 CDK1, PLK1, cdc25c, and CHK1, and increased p16, p21, and p27), likely as a consequence of RAS/MAPK signaling inhibition (**Figure 4B**). Further, there was a marked shift to an epithelial phenotype, as indicated by increased EMA, EPPK1, Claudin1, and beta-catenin (**Figure 4B**). Many of the associations with sensitivity to cobimetinib were identifiable in the pre-treatment samples, with the associations markedly accentuated and extended in cobimetinib-treated samples. The marked increase in BIM in response to MEKi has been identified previously and provides a biomarker for response to combined inhibition of MEKi and BCL2 family members (Cragg et al., 2008; Iavarone et al., 2019). We also performed a similar analysis using trametinib (**Figure S3**) and observed a marked overlap of potential biomarkers despite the analysis of different cell lines and different MEK inhibitors. Importantly, the key adaptive pathway-level changes associated with drug sensitivity include cell cycle inhibition in sensitive cell lines (t-test, p = 4.1 ×10^−4^, **Figure 4C**) and PI3K/Akt signaling activation in resistant cell lines (t-test, p = 0.015, **Figure 4C**). Together, the results argue that (i) sensitivity to RAS/MAPK pathway inhibition is associated with baseline pathway activity and cell state, and (ii) adaptive responses to RAS/MAPK pathway inhibition in resistant cells could be overcome by PI3K inhibitors (although the toxicity of combinations of RAS/MAPK and PI3K pathway inhibitors has to be considered).

### A systematic “protein-drug” connectivity map

To systematically evaluate the utility of our protein response data, we built a protein-drug connectivity map based on the RPPA data. In this map, each node represents a protein or a drug, protein-drug connections are based on whether the drug treatment caused a significant change of the protein, and drug-drug connections are based on whether the two drugs caused similar protein responses (**Figure 5A**). As expected, drugs for the same target are clustered together: for example, several MEK inhibitors and mTOR/PI3K inhibitors are highly connected, highlighting their similar downstream protein responses. This map also identifies novel connections: a PARP inhibitor showed both similar and opposite relationships with some drugs, suggesting potential additive or agonistic effects that could direct the development of rational drug combinations. Indeed, based on assessment of functional proteomics changes as assessed by RPPA, we have validated synergistic activity of PARP inhibitors and inhibition of PI3K pathway, MEK, ATR, and WEE1 inhibitors in preclinical and clinical studies (Fang et al., 2019; Shen et al., 2015; Sun et al., 2017; Sun et al., 2018).

**Figure 5.**
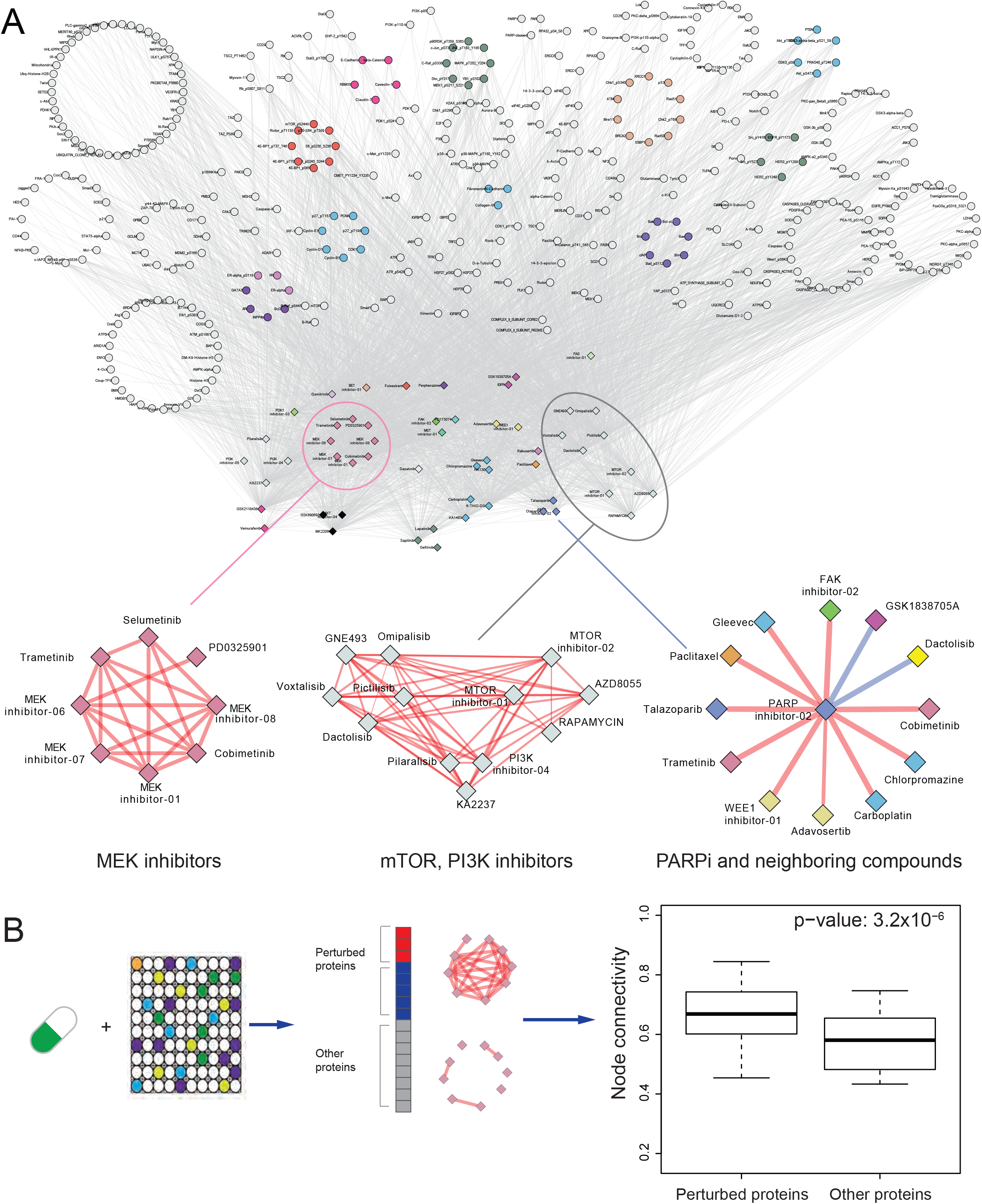
A “drug-protein” connectivity map based on protein response signals. (A) A global view of the drug-protein connectivity map with highlighted examples of drug-drug correlation networks (i.e., MEK inhibitors, mTOR, PI3K inhibitors, and neighboring drugs of a PARP inhibitor). Red/blue edges represent positive/negative drug-drug correlations, respectively. Proteins were grouped and colored by their related functional pathways. The drugs were grouped and colored by their targeted genes or pathways. (B) Comparison of node connectivity between perturbed and neutral proteins in the protein interaction network. P-value was computed based on a paired Wilcoxon test.

We next studied protein-protein relationships in the map. For any given drug treatment, we classified proteins into perturbed proteins and other proteins. We found that perturbed proteins are more likely to interact than other proteins, based on the STRING database (Szklarczyk et al., 2019) (t-test, p = 3.2 ×10^−6^), suggesting that proteins co-perturbed by a drug tend to be involved in the same biological processes and to interact as part of a signaling cascade (**Figure 5B**). This global assessment using prior protein interaction knowledge supports the utility of the approach to drive biological discoveries.

Using drug-centered protein neighborhoods, we initially focused on signaling through tyrosine kinases and their downstream networks: selumetinib (target: MEK) (**Figure 6A**), AZD8055 (target: mTOR) (**Figure 6B**), GSK1838705A (target: IGF1R/ALK) (**Figure 6C**), and sapatinib (target: EGFR/ERBB2) (**Figure 6D**), and demonstrated a marked overlap in protein networks in inhibitor-perturbed cells. Interestingly, the Hsp90 inhibitor (gamitrinib) protein neighborhood (**Figure S4A**), demonstrated similarities to that of the tyrosine kinase pathway inhibitors, potentially due to a role of Hsp90 in stabilizing multiple members of the tyrosine kinase signaling pathway. Indeed, the similarities in the protein networks argue that the major effects of Hsp90 are likely attributable to its effects on tyrosine kinase signaling pathways (Lee et al., 2017). In contrast, rabusertib (target: Chk1) (**Figure S4B**), and chlorpromazine (target: autophagy) (**Figure S4C**) demonstrated distinct protein neighborhoods consistent with markedly different mechanisms of action.

**Figure 6.**
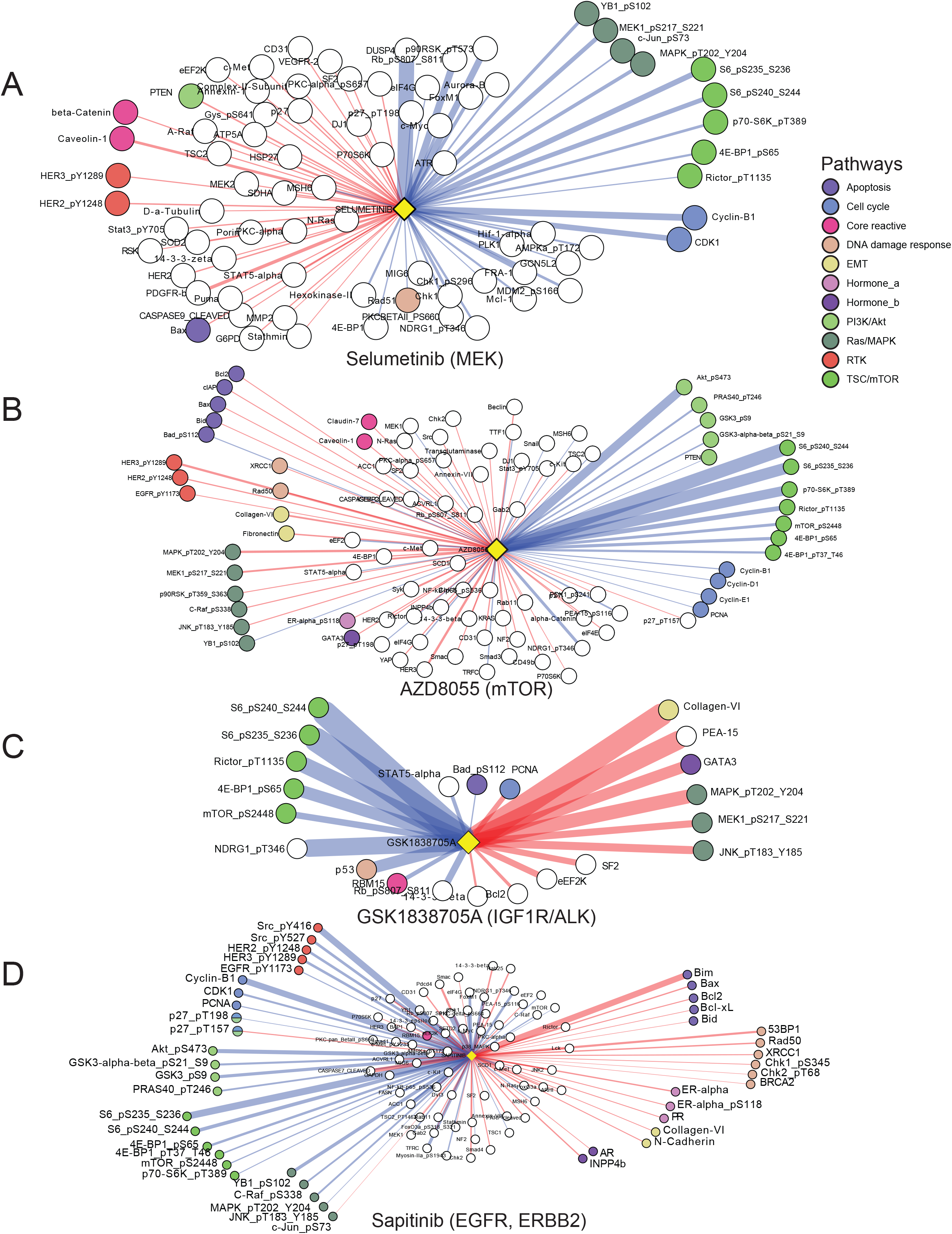
Examples of drug-centered protein connectivity maps. (A-D) Drug-protein connectivity maps for individual drugs: selumetinib (A), AZD8055 (B), GSK1838705A (C), and sapitinib (D). In the network view, the edge color indicates the response direction of protein markers (red/blue: up/down-regulated in post-treatment). The proteins (nodes) of the same functional pathways are highlighted in the same colors. See also Figure S4.

As described above, the MEKi protein neighborhood is strongly associated with signaling through the MAPK and mTOR pathways, cell cycle progression, and cell state. There is also a strong association with apoptotic balance (BIM, BAX, and MCL1). Based on extensive validation of the relationships between these pathways and RAS/MAPK signaling, the association with multiple other proteins in the neighborhood map in **Figure 6A** are likely valid. Given that the MAPK pathway is a key regulator of the TSC1/2 complex that is upstream of mTORC1 signaling, it is not surprising that the protein neighborhood of mTOR inhibitor AZD8055 (**Figure 6B**) is highly related to the selumetinib protein neighborhood. The most marked differences between the MEK and mTOR inhibitor protein neighborhoods are represented in the upper components of the PI3K and MAPK pathway that appear relatively independent of each other. Interestingly, the IGF1R/ALK inhibitor, GSK1838705A, protein neighborhood encompasses components of both the MEK and mTOR protein neighborhoods, consistent with the IGF1R having input into both pathways. While the strong link to the PI3K pathway was expected, the link between the IGF1R and MAPK pathway has been less studied. The pan-EGFR family inhibitor, sapatinib, neighborhood reflects EGFR family receptors being the key regulators of the PI3K and MAPK pathways in epithelial cells (Akbani et al., 2014). The EGFR family has a stronger link than either mTOR or MEK inhibitors to the DNA damage repair pathway (i.e., 53BP1, Rad50, XRCC1, pChk1/2, and BRCA2), consistent with recent studies (Russo et al., 2019; Wang et al., 2013).

### A user-friendly data portal for protein responses of perturbed cell lines

To facilitate the utilization of our protein response data by a broad biomedical community, we provided unrestricted access to the data through a user-friendly portal, called “Cancer Perturbed Proteomics Atlas” for fluent data exploration and analysis, which can be accessed at http://bioinformatics.mdanderson.org/main/:CPPAOverview. The data portal provides four interactive modules: “Data Summary,” “My Protein,” “Connectivity Map,” and “Analysis” (**Figure 7i**). The “Data Summary” module provides detailed information about each sample (including cell line, compound, dose, time, and culture conditions) (**Figure 7ii**). The datasets can be easily downloaded through a tree-view interface. “My protein” module provides annotation of RPPA protein markers, including the corresponding genes, and antibody information (**Figure 7iii**). The “Connectivity Map” provides an interactive approach to exploring the map, through which protein-drug and drug-drug connectivity can be examined through different visual and layout styles (**Figure 7iv**). The “Analysis” module provides three common analyses through which users can explore protein responses associated with a drug/compound, including protein response (Δp) rank (**Figure 7v**), volcano plots for the correlations between protein responses and drug sensitivity (**Figure 7vi**), and box plots for differential protein responses between sensitive and resistant cell lines (**Figure 7vii**). Collectively, this effort provides a valuable platform that enables researchers to explore, analyze, and visualize RPPA-based protein response data intuitively and efficiently.

**Figure 7.**
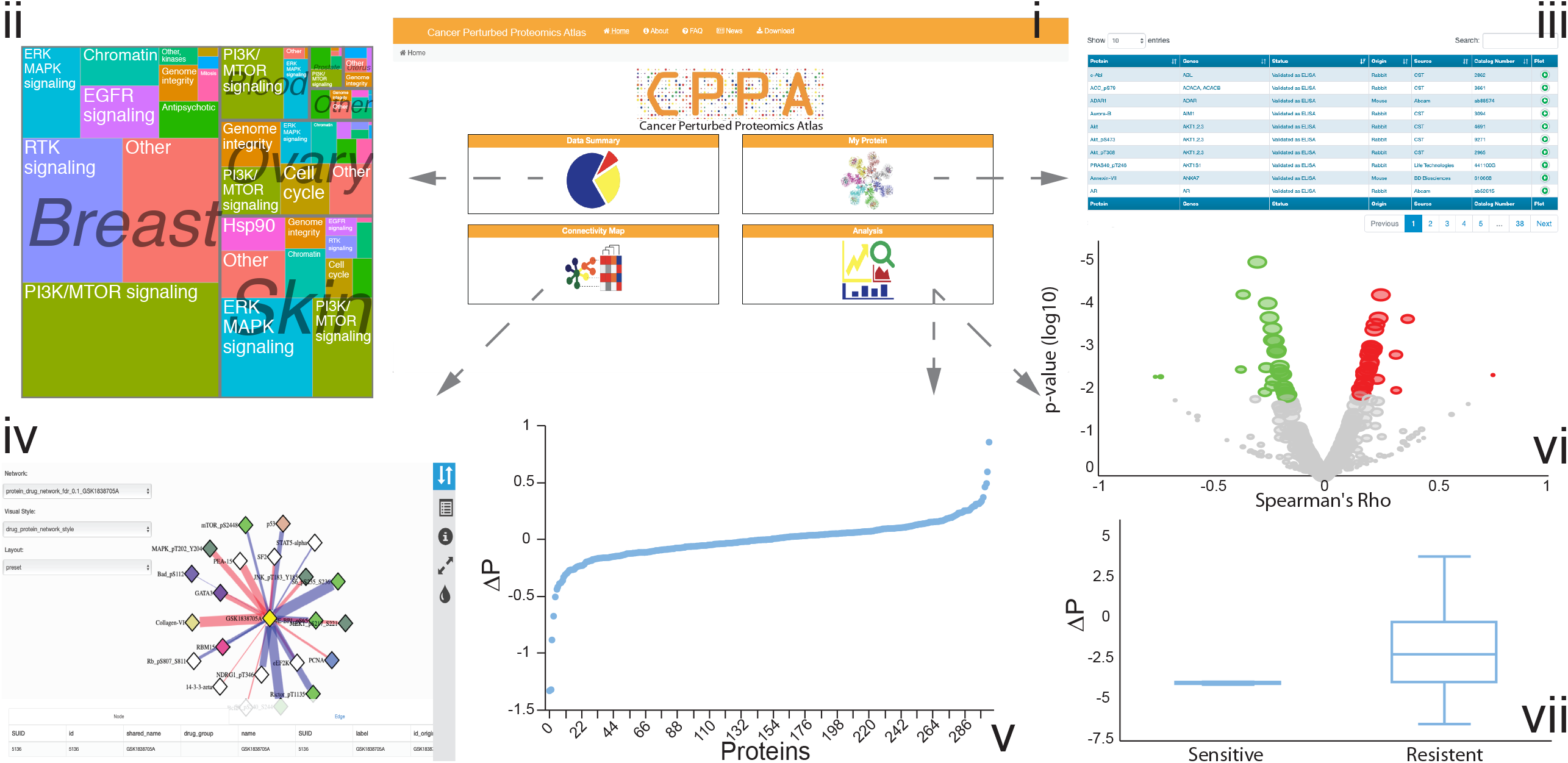
A snapshot of the data portal for the perturbed RPPA data. The CPPA portal (i) contains four modules: the “Summary” module (ii); the “My protein” module (iii); the “Network Visualization” module (iv); and the “Analysis” module, which offers protein perturbation analysis (v), correlation analysis of drug and protein response (vi), and differential expression analysis of protein response in sensitive and resistant cell lines (vii).

## Discussion

Understanding the functional consequences of drug treatment is central to identifying patients who will likely benefit from specific therapies and to develop effective personalized combination cancer therapies. Here we present a large collection of protein responses (including total and post-translationally modified proteins) upon drug treatments (>12,000 treated samples) using RPPA, the same platform employed for protein profiling of TCGA patient samples and CCLE cell line samples. Our dataset is several magnitudes larger than previously published studies, and for such a resource, data quality is key. We validated the quality of our datasets in several ways. First, we demonstrated the high reproducibility of technical replicate samples using the same platform. Second, we established a high consistency between RPPA-based protein responses and the independently generated L1000 mRNA responses to the same perturbation conditions. Third, we validated observed correlations of protein responses with public drug sensitivity data using newly generated drug sensitivity data on independent cell line sets. Finally, the quality of our dataset is also supported by the meaningful patterns observed on the systematic “protein-drug” connectivity map, such as the clustering of similar drugs and higher node connectivity of perturbed proteins annotated in the STRING protein interaction database. Our study represents a unique, high-quality compendium of protein responses of cancer cell lines to a diversity of compound perturbations available for use by the wider community.

By establishing the connections between a core set of proteins and drug treatment in simplified well-controlled cell line models, the utility of our protein response dataset is several-fold. First, our dataset provides a basis for understanding cause-effect relationships, which is complementary to correlation analyses and associations that can be obtained from patient cohorts. Based on these data, it will be possible to develop quantitative predictive models of how signaling networks function in intact cellular systems. Second, we show that while there is information content in biomarkers at baseline, the information content is markedly increased when baseline and response signals are combined. This is predicted by systems biology and engineering precepts, wherein, perturbed systems contain more information than static analysis. Biomarkers designed to select treatment using baseline data frequently have a limited power to predict benefit, and our results suggest that adaptive protein responses after initial treatment could be highly informative in terms of treatment response, and clinical benefit. Further clinical investigations are warranted to assess the potential benefit gains using such a strategy. Third, since protein responses reflect how cancer cells critically rewire their signaling pathways to survive and adapt to the stress of a specific drug treatment, these protein signals provide a strong basis for rational design of combination therapies as we have demonstrated previously (Iavarone et al., 2019; Krepler et al., 2017; Krepler et al., 2016; Kwong et al., 2015; Molinelli et al., 2013; Muranen et al., 2012; Sun et al., 2017; Sun et al., 2018).

We recognize some limitations of this study. First, compared with mass spectrometry-based protein level or mRNA level readout assays, the number of protein markers that can be effectively monitored by the RPPA technology is much smaller. However, the increased sensitivity (particularly for some key proteins and phosphoproteins), and cost considerations, make RPPA a practical platform for generating such a large resource. In capturing protein responses, RPPA and mass spectrometry are complementary because of their different scopes and focuses. Second, although many perturbed protein response profiles were generated, some cell lines and drug treatments (including different dosages) are still sparsely sampled. Further efforts are required to obtain more compensative sets; moreover, machine learning approaches may have the potential to fill some of these gaps. Finally, as with other high-throughput technologies, there can be technical measurement errors for individual samples, and interesting observations from our study should be followed by further in-depth investigations.

We have provided an interactive, user-friendly data portal through which biomedical researchers can explore, visualize, and intuitively analyze these data. With this bioinformatics tool, we expect an effective translation of the large-scale perturbed protein data into biological knowledge and clinical utility. Together with recent efforts that have systematically characterized phenotypic and molecular responses to drug treatment, our study provides a rich resource for the research community to investigate the behaviors of cancer cells and the dependencies of treatment responses.

## Acknowledgements

This study was supported by the National Institutes of Health (U01CA168394, U24CA143883, U54HG008100, P50CA098258, and P50CA217685 to G.B.M., U24CA209851 and U01CA217842 to H.L. and G.B.M., U24CA210950 and U24CA210949 to R.A. and G.B.M., and Cancer Center Support Grant P30CA016672), a kind gift from the Sheldon and Miriam Adelson Medical Research Foundation, Susan G Komen (SAC110052), Ovarian Cancer Research Foundation (545152), and Breast Cancer Research Foundation (BCRF-18-110) to G.B.M., a DoD/CDMRP (W81XWH-16-1-0237) to R.A., and the Lorraine Dell Program in Bioinformatics for Personalization of Cancer Medicine (to H.L.). We thank Kamalika Mojumdar for editorial assistance. RA is a bioinformatics consultant for the University of Houston. G.B.M. is on the Scientific Advisory Board or a consultant for AstraZeneca, ImmunoMet, Lilly, Nuevolution, PDX Pharmaceuticals, Symphogen, and Tarveda, has stock options with Catena Pharmaceuticals, Immunomet, Signalchem, and Tarveda, and has licensed technology to Myriad and Nanostring. H.L. is a shareholder and scientific advisor of Precision Scientific Ltd.

## Author Contributions

G.B.M. and H.L. conceived of the project. W.Z., J.L., M.C., J.Z., A.S., S.V.K., A.K., R.A., G.B.M., and H.L. contributed to the data analysis. N.K.N., K.J.C., C.T.B, Y. Lawrence, N.T.P., M.A.D., M.H., T.M., I.Z., E.V.E., P.T.S., D.J.S., J.S.B., J.W.G., Y. Lu and G.B.M. contributed to the experiments. M.C. and J.L. implemented the web portal. W.Z., J.L., M.C., G.B.M., and H.L. wrote the manuscript, with input from other authors. H.L. supervised the whole project.

## STAR methods

### Contact for Reagent and Resource Sharing

Further information and requests for resources and reagents should be directed to and will be fulfilled by the Lead Contact, Han Liang.

### Experimental Model and Subject Details

#### Cell lines

We collected cancer cell lines through the MD Anderson Cancer Center (MDACC) CCSG supported Cell Line Characterization Core Facility (Houston, TX, USA) and from several outside collaborations. All cell lines prepared at MDACC were confirmed by short tandem repeat (STR) analysis in the core per institutional policy, and the outside collaborators also routinely confirmed cell lines by STR analysis.

#### RPPA experiments

Cell line samples were prepared, and antibodies were validated as previously described (Hennessy et al., 2010; Li et al., 2017). RPPA data were generated by the RPPA core facility at MDACC. RPPA slides were first quantified using ArrayPro (Media Cybernetics) to generate signal intensities (level 1), then processed by SuperCurve to estimate the relative protein expression level (level 2), and were then normalized by median polish (level 3). RPPA slide quality was assessed by a quality control classifier (Ju et al., 2015), and only slides above 0.8 (range: 0-1) were retained for further analysis. In total, we generated RPPA data from 15,867 samples, including 12,222 treated cell line samples and 3,647 baseline samples (e.g., treated by DSMO) from 8 batches. Through the comparison of RPPA signals between post-treatment (p1) and baseline samples (p0), we generated 8,111 unique protein response (Δp) profiles after combining replicate samples.

#### Comparison of RPPA-based protein response and L1000-based mRNA response

To validate our RPPA perturbation data, we downloaded the level-5 data of L1000 phase 1 from the GEO database (GSE92742). For a fair comparison, we collected data from the same cell lines perturbed by the same compound. In total, 46 “perturbation-cell-line” IDs (60 samples) and 316 genes/proteins (total proteins only) commonly shared by the two platforms were used in the subsequent analyses. For a perturbation-cell-line ID with multiple concentration and/or time points, we adopted the median value across all conditions as the representative response score. As shown in **Figure 2A**, for each platform, we first converted the continuous response to a categorical response: up-regulated, down-regulated, or neutral. Random events were defined by the global median ± 35% quantile, calculated from the full matrix. Next, we excluded the random events and computed Goodman-Kruskal’s gamma to estimate sample associations across genes. We evaluated the concordance between RPPA and L1000 platforms through two analyses. (i) Protein-mRNA response associations: for each sample, a gamma (ɣ) association between the two platforms was computed across genes/proteins when at least 12 genes showed up-/down-regulation. To generate the background distribution, we randomly shuffled protein labels and computed the response associations between the shuffled proteins and mRNAs (the seed used for randomization is “1234”). Then, a paired Student’s t-test was used to evaluate the statistical significance of the group difference between the real and matched randomly shuffled responses. (ii) Sample-sample associations: in our RPPA dataset, a perturbation-cell-line ID might have replicate samples. Here, we only retained the one with the best protein-mRNA response association from the previous analysis. Next, for samples that showed up-/down-regulation of >3 genes, gamma associations for every two such samples were computed within each platform (within the same batch). Then, Pearson’s correlation between the significant gramma associations (FDR < 0.01 for each platform) was used to evaluate the consistency between the protein and mRNA responses.

#### Analysis of predictive protein markers of drug sensitivity

We collected drug sensitivity data from two databases: GDSC and CTRPv2. For validation, an in-house drug screening data set was generated for selected compounds. RPPA perturbation data of the same cell line treated with the same compound at different dosages or time points were averaged using mean values. The baseline level (p0) and protein response levels (Δp) were tested for associations with drug sensitivity (IC50 or AUC score) in univariate linear models. The joint markers (Δp|p0) were defined as the predictions of linear regression models, including both baseline and protein response for specific antibodies. Predictive markers were selected at a significance level of p = 0.05. To identify the differential protein markers of drug sensitivity and resistance, cell lines were classified as sensitive or resistant to a specific drug based on the consensus call of CTRPv2, GDSC, and in-house datasets. Baseline levels (p0) and protein response levels (Δp) with a significant difference between sensitive and resistant cell lines were identified by Student’s t-test. Pathway-level scores (Li et al., 2017) were similarly analyzed.

#### Construction of a drug-protein connectivity map

The association of each drug-protein pair was assessed by testing the difference of protein expression between baseline (p0) and post-treatment level (p1) based on the paired t-test across cell lines. For each drug-drug pair, we used Goodman-Kruskal’s gamma to calculate the associations, as described in the comparison between RPPA-based protein response and L1000-based mRNA response data. The significantly correlated drug-protein and drug-drug pairs (FDR < 0.1) were used to construct a global drug-protein connectivity map. In the connectivity map, proteins were grouped and colored by their related protein functional pathways, and drugs were grouped and colored by their targeted genes or pathways. For each drug, the network densities were calculated for the two subsets of proteins: (i) proteins significantly differentially expressed between p0 and p1 (perturbed proteins), and (ii) other proteins (neutral proteins). The network density *D* of a protein subset with size *N* was defined as a ratio of the number of protein-protein interactions (*E*) to the number of all possible protein pairs 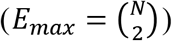, i.e., *D* = *E*/*E*_*max*_. Protein-protein interaction information was obtained from the STRING database (Szklarczyk et al., 2019). A paired Wilcoxon test was performed to assess the difference of the network densities between the perturbed and other proteins of all the drugs. For **Figure 6**, the examples of drug-centered connectivity maps were generated separately with colored edges (red: up-regulated in post-treatment; blue: down-regulated in post-treatment). The edge widths were proportional to the differential expression between baseline (p0) and post-treatment levels (p1). All network views were generated by the Rcy3 library and Cytoscape (Otasek et al., 2019; Shannon et al., 2003).

#### Data portal development

All RPPA, mRNA expression, and drug sensitivity data accompanying the pre-calculated analytic results were stored in a CouchDB database. We generated all the analytic results in R before loading them into the database. We implemented a user-friendly and interactive web interface in JavaScript. Specifically, tabular results were generated by DataTables, box, and scatter plots were generated by HighCharts, and interactive network views were implemented by Cytoscape.js library.

## Supplementary Figures

**Figure S1. Summary of the perturbed RPPA profiling data in this study, related to Figure 1.**

(A) Cell lines associated with the largest sample sizes. (B) Sample numbers by different types of compound perturbations. (C) Compounds associated with the largest sample sizes.

**Figure S2. Confirmation of predictive protein markers for selected drugs, related to Figure 3**

(A-C) Pictilisib (drug response data from CTRPv2) and (D) talazoparib (drug response data from in-house experiments). The analysis was based on drug sensitivity data independent from Figure 3. (A, D) The heatmaps show Pearson’s correlation coefficients of protein level and drug sensitivity. The bar plots show the number of predictive protein markers in each category. The protein markers of baseline (p0) and protein response (Δp) were selected at the significance level of p = 0.05 in the univariate correlation test. The joint protein markers, labeled as “Δp|p0,” are the linear regression models of baseline and protein response levels for specific proteins. The coefficients in the heatmap for joint markers indicate the correlation between the prediction of the linear model and the drug sensitivity. Δp total is defined as the union of “Δp only” and “Δp given p0.” (B, C) The scatter plots summarize the comparison of the Pearson’s correlation coefficients of drug sensitivity and (B) baseline level (p0) as well as (C) protein response (Δp) in the two independent data sets.

**Figure S3. Differentially expressed proteins between trametinib-sensitive and -resistant cell lines, related to Figure 4**

(A, B) Heatmaps showing baseline protein expression (A), and perturbed protein response (B) with a significant difference between the sensitive and resistant cell lines (q < 0.05).

**Figure S4. Examples of drug-centered protein connectivity maps, related to Figure 6**

(A-C) Drug-protein connectivity maps for individual drugs: gamitrinib (A), rabusertib (B), and chlorpromazine (C). In the network views, the edge color indicates the response direction of protein markers (red/blue: up/down-regulated in post-treatment). The proteins (nodes) of the same functional pathways are highlighted in the same colors.

