## Supplementary figures for "Large-scale Characterization of Drug Responses of Clinically Relevant Proteins in Cancer Cell Lines"

### Figure S1

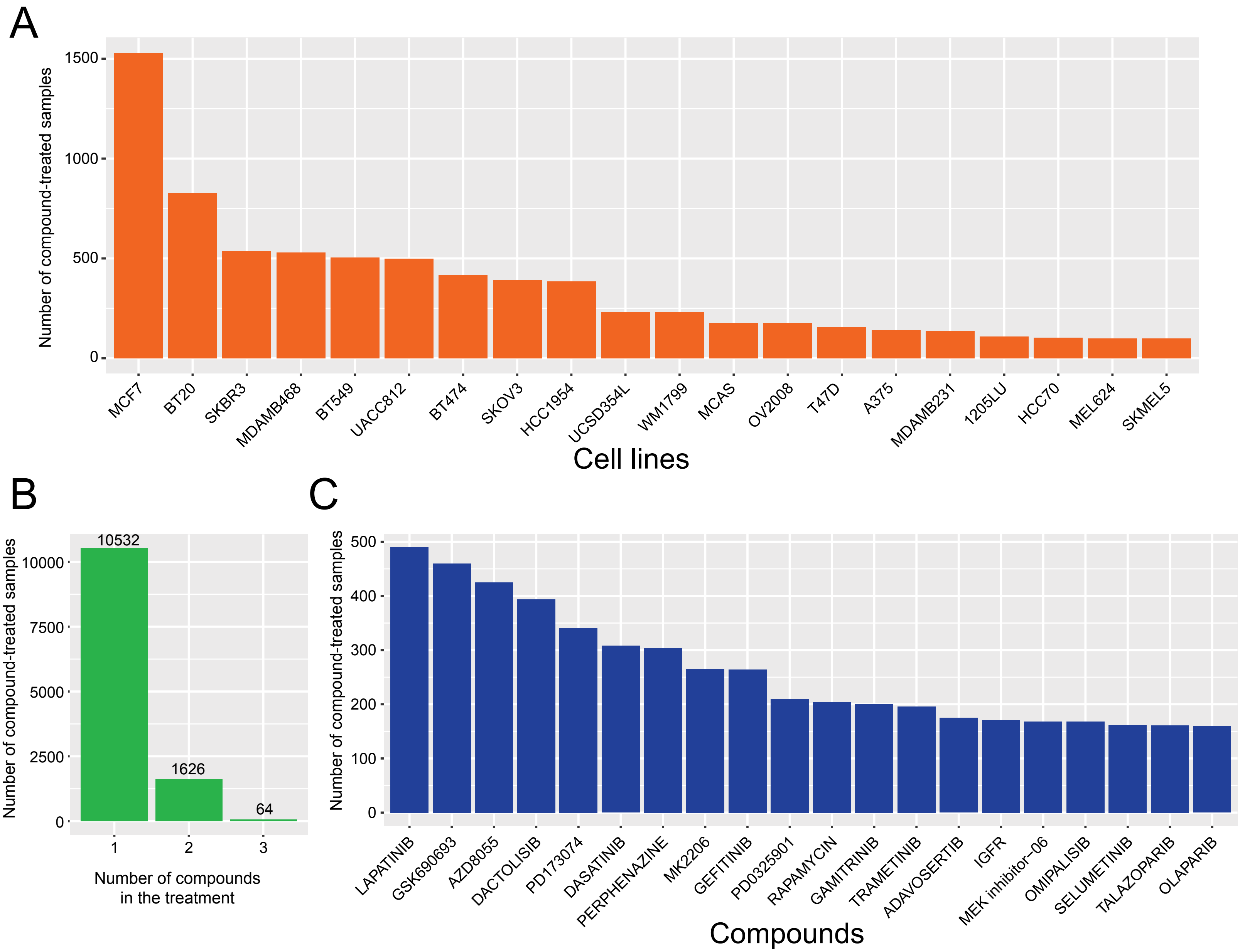


**Figure S1. Summary of the perturbed RPPA profiling data in this study, related to Figure 1.**
(A) Cell lines associated with the largest sample sizes. (B) Sample numbers by different types of compound perturbation. (C) Compounds associated with the largest sample sizes.

### Figure S2

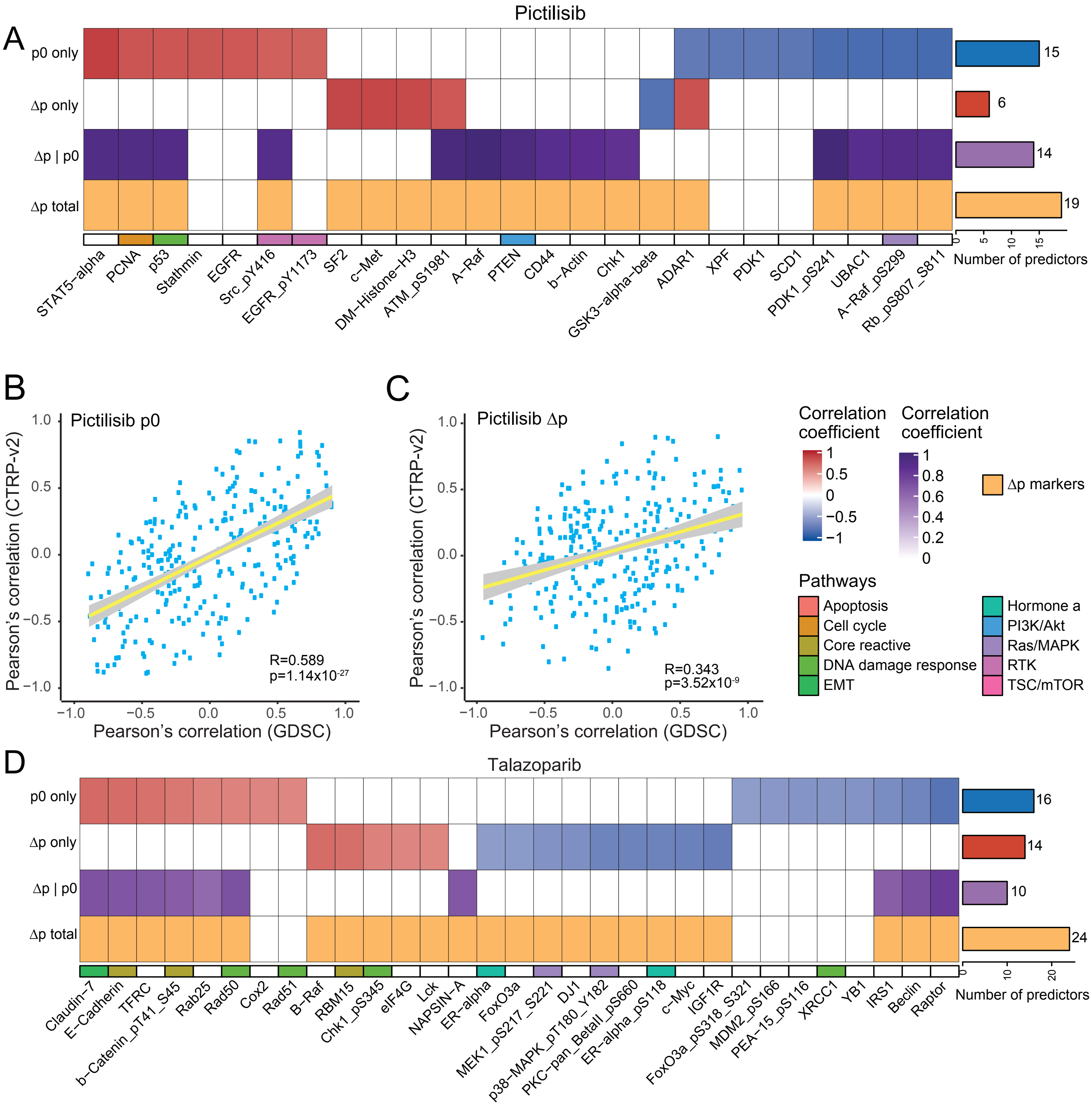


**Figure S2. Confirmation of predictive protein markers for selected drugs, related to Figure 3.**

(A-C) Pictilisib (drug response data from CTRPv2), (D) talazoparib (drug response data from in-house experiments). The analysis was based on drug sensitivity data independent from Figure 3. (A, D) The heatmaps show Pearson's correlation coefficients of protein level and drug sensitivity. The bar plots show the number of predictive protein markers in each category. The protein markers of baseline (p0) and protein response (Δp) were selected at the significance level of p = 0.05 in the univariate correlation test. The joint protein markers, labeled as "Δp|p0," are the linear regression models of baseline and protein response levels for specific proteins. The coefficients in the heatmap for joint markers indicate the correlation between the prediction of the linear model and the drug sensitivity. Δp total are defined as the union of "Δp only" and "Δp given p0." (B, C) The scatter plots summarize the compari-son of the Pearson's correlation coefficients of drug sensitivity and (B) baseline level (p0) as well as (C) protein response (Δp) in the two independent data sets.

### Figure S3

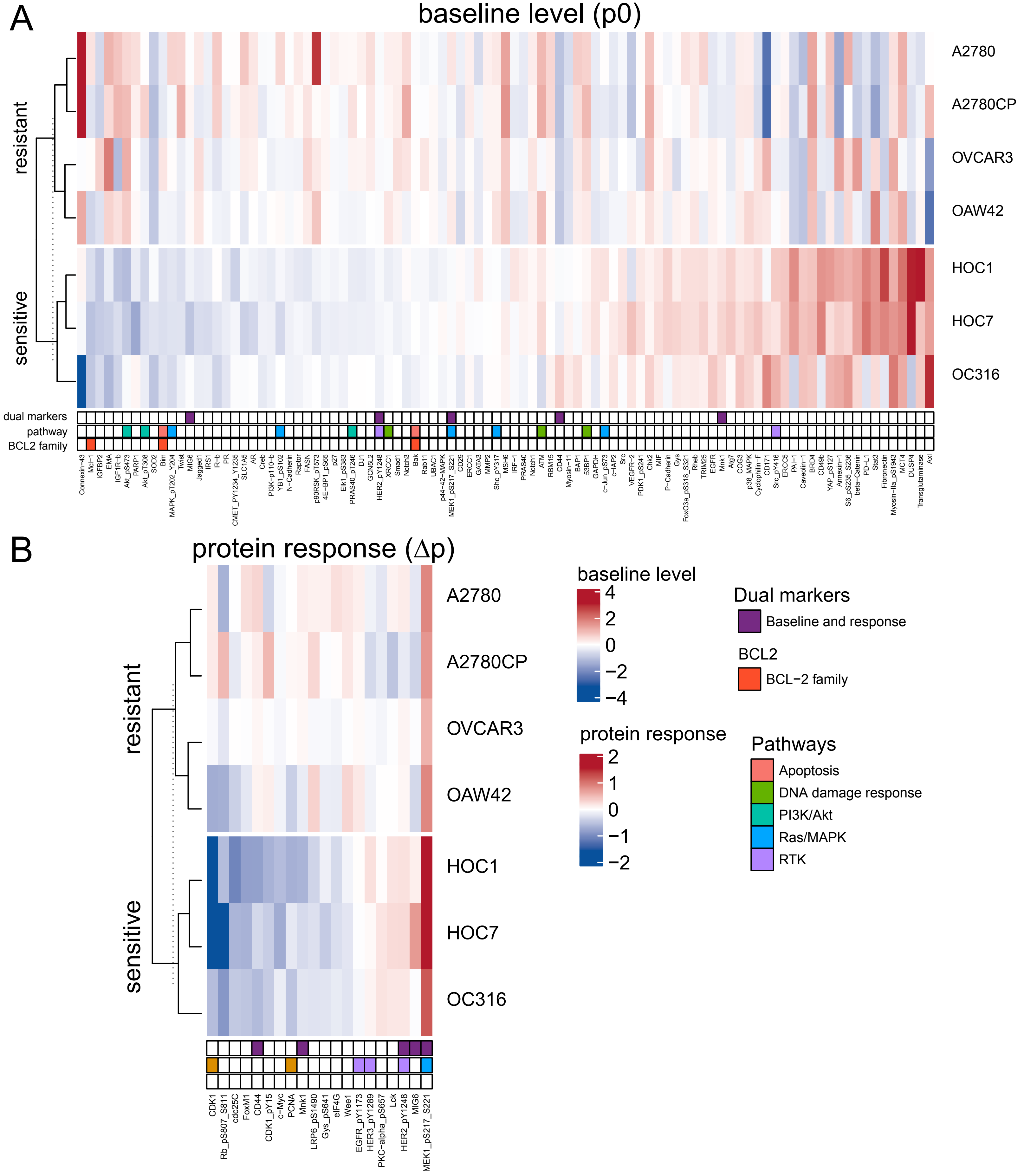


**Figure S3. Differentially expressed proteins between trametinib-sensitive and -resistant cell lines, related to Figure 4.**

(A, B) Heatmaps showing baseline protein expression (A), and perturbed protein response (B) with a significant difference between the sensitive and resistant cell lines ($q < 0.05$).

### Figure S4

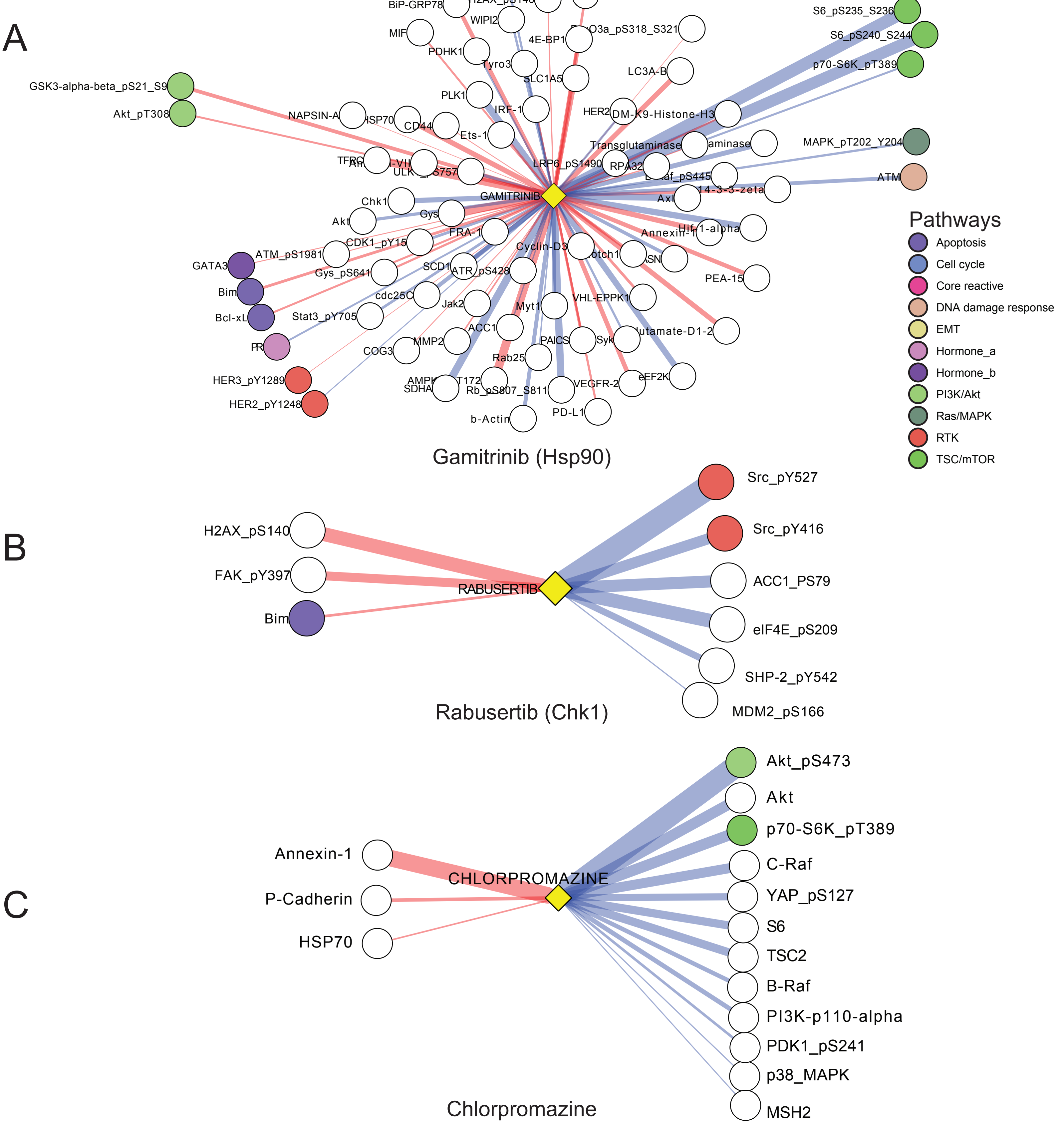


**Figure S4. Examples of drug-centered protein connectivity maps, related to Figure 6.**
(A-C) Drug-protein connectivity maps for individual drugs: gamitrinib (A), rabusertib (B), and chlorpromazine (C). In the network views, the edge color indicates the response direction of protein markers (red/blue: up/down-regulated in post-treatment). The proteins (nodes) of the same functional pathways are highlighted in the same colors.
